# Small-molecule antibiotic inhibitors of post-translational protein secretion

**DOI:** 10.1101/2020.11.17.387027

**Authors:** Mohamed Belal Hamed, Ewa Burchacka, Liselotte Angus, Arnaud Marchand, Jozefien De Geyter, Maria S. Loos, Jozef Anné, Hugo Klaassen, Patrick Chaltin, Spyridoula Karamanou, Anastassios Economou

## Abstract

The increasing problem of bacterial resistance to antibiotics underscores the urgent need for new antibacterials. The Sec preprotein export pathway is an attractive potential alternative target. It is essential for bacterial viability and includes components that are absent from eukaryotes. Here we used a new high throughput *in vivo* screen based on the secretion and activity of alkaline phosphatase (PhoA), a Sec-dependent secreted enzyme that becomes active in the periplasm. The assay was optimized for a luminescence-based substrate and was used to screen a ~240K small molecule compound library. After hit confirmation and analoging, fourteen HTS secretion inhibitors (HSI), belonging to 8 structural classes, were identified (IC_50_ <60 μM). The inhibitors were also evaluated as antibacterials against 19 Gram^−^ and Gram^+^ bacterial species (including those from the WHO top pathogens list). Seven of them, HSI#6, 9; HSI#1, 5, 10 and HSI#12, 14 representing three structural families were microbicidals. HSI#6 was the most potent (IC_50_ of 0.4-8.7 μM), against 13 species of both Gram^−^ and Gram^+^ bacteria. HSI#1, 5, 9 and 10 inhibited viability of Gram^+^ bacteria with IC_50_ ~6.9-77.8 μM. HSI#9, 12 and 14 inhibited viability of *E. coli* strains with IC_50_ <65 μM. Moreover, HSI#1, 5 and 10 inhibited viability of an *E. coli* strain missing TolC to improve permeability with IC_50_ 4-14 μM, indicating their inability to penetrate the outer membrane. *In vitro* assays revealed that antimicrobial activity was not related to inhibition of the SecA component of the translocase and hence HSI molecules may target new unknown components that affect secretion. The results provide proof of principle for our approach, and new starting compounds for optimization.

## Introduction

Development of modern medical diagnostics and theapeutics, vaccination programs and improved living standards have led to the control and even elimination of many infectious diseases (Rodriguez-Rojas et al., 2013). Antibiotics have been major contributors to this outcome and one of the most important discoveries of the pharmaceutical industry of the 20th century (Davies and Davies, 2010). However, their use is currently limited due to the increasing antibiotic resistance of various bacterial strains and to undesirable side effects (Rodriguez-Rojas et al., 2013). Antimicrobial resistance is responsible for an estimated 25,000 deaths and 1.5 billion € in healthcare costs/year in the European Union (Comission, 2015). Therefore, there is an urgent need to develop new strategies and methods to prevent epidemics.

Finding new antibiotics against new bacterial target proteins is challenging. Novel antibiotic targets should be: (i). essential for bacterial growth. (ii). ideally, conserved in bacteria, so that antibiotics can have a broad spectrum, but not present in eukaryotes or, if present, should be sufficiently diverged or inaccessible. A parallel approach for new anti-infectives are drugs that inhibit bacterial virulence and/or pathogenesis but are not essential for viability. Such inhibitors can be part of combination therapies and can boost effective immune responses in the host (Calvert et al., 2018).

One important process for both bacterial viability and virulence is protein secretion into and across the plasma membrane using the Sec system (Fig. 1A) (Tsirigotaki et al., 2017; Tsirigotaki et al., 2018). It mediates the export of 30-35% of the bacterial proteome and of >90% of bacterial effectors and toxins (Driessen and Nouwen, 2008) and includes 18 components essential for viability (Goodall et al., 2018; Loos et al., 2019) and dozens essential for virulence. Sec secretion pathway components like the ATPase SecA (Rao et al., 2014), the BAM outer membrane assembly complex (Kim et al., 2012), signal peptidases(Rao et al., 2014) and the lipoprotein trafficking system LOL (Zuckert, 2014) meet wholly or partly the criteria of attractive targets. Many of these proteins have important advantages as drug targets: they are stable *in vitro*, have rather well known structure/function features and probably good accessibility due to their location in the cell envelope and membranes (Rao et al., 2014). These export machineries are then used by 59 proteins that are essential for viability that are exported in K-12 MG1655 *E. coli* and are located in the inner membrane and periplasm (Loos et al., 2019).

**Figure 1.**
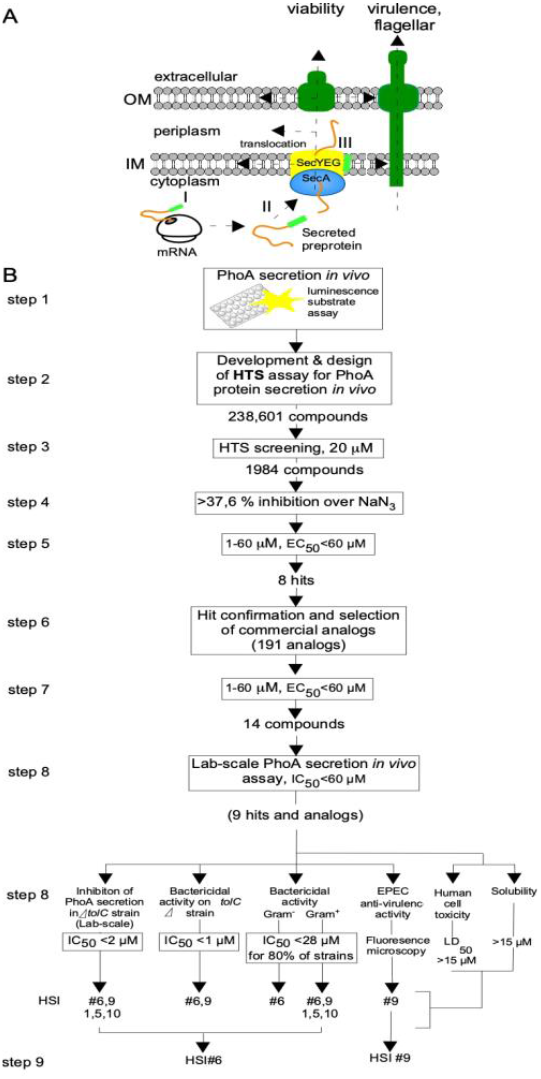
Overview of Sec pathway and HTS pipeline to discover secretion inhibitors. **A.** Cartoon of the Sec pathway in a cell. **B.** HTS and screening pipeline used for the identification and characterization of secretion inhibitors.

In the Sec secretion system, SecA converts ATP energy to drive the translocation of preproteins across the SecY, SecE and SecG channel in the bacterial cytoplasmic membrane. Exported preproteins containing *N*-terminal signal sequences transit the cytoplasm in non-folded states, with or without bound chaperones (Kusters and Driessen, 2011; Chatzi et al., 2014; Busche et al., 2018; Hamed et al., 2018b). After preprotein translocation into the periplasm (Gram^−^) or extracellular space (Gram^+^), signal sequences are removed by membrane-embedded signal peptidases and exported proteins acquire their native structures or are further trafficked (Kusters and Driessen, 2011; Hamed et al., 2018a).

Apart from the generalized Sec system, several specialized mechanisms control the translocation of pathogenicity proteins to the bacterial cell surfaces and beyond (Holland, 2004). Invariably, assembly of all these transport machineries is Sec-dependent wholly or partially and trans-envelope “conduits” formed usually take toxins directly to the outside milieu or even to host cells (Stathopoulos et al., 2000; Natale et al., 2008). One Sec-independent protein secretion virulence mechanism in Gram^−^ bacteria is the type III secretion system (T3SS) that injects toxins directly in host cell cytosols (Galan and Collmer, 1999; Portaliou et al., 2016; Deng et al., 2017). During infection, enteropathogenic EPEC *E.coli* cells use their T3SS to attach to epithelial cells, colonize them and induce the formation of actin pedestals (Goosney et al., 2000; Shaw et al., 2001).

For targets like SecA that have several possible sites of action for inhibition, there are potentially several assays that can be used. *In vitro* assays include measuring translocation ATPase, seen upon SecA binding to both membranes and preprotein, commonly by following released phosphate using the malachite green reagent (Sugie et al., 2002; Dietrich et al., 2010; Gouridis et al., 2010). *In vivo* assays of protein secretion can be based in whole cell assays (Alksne et al., 2000), and monitor reduced expression of SecA (Parish et al., 2009) or cytoplasmic accumulation of β-galactosidase if translocation is blocked (Crowther et al., 2012).

Several SecA inhibitors have been discovered using different screens (Oliver et al., 1990; Rao et al., 2014). Sodium azide, the first known SecA inhibitor (Oliver et al., 1990) is not a usable antibacterial because it is non-selective and inhibits eukaryotic enzymes such as other ATPases including mitochondrial F-ATPase (Bowler et al., 2006), cytochrome c oxidase, superoxidase mutase and alcohol dehydrogenase (Li et al., 2008). Equisetin (CJ-21,058) inhibits translocation ATPase (IC_50_ of 15 μg/ml) (Sugie et al., 2002). 5-amino-thiazolo(4,5-D)pyrimidine revealed low IC_50_ values (>135 μM) against translocation ATPase of *E.coli* SecA (Segers and Anne, 2011). Hydroxanthene derivatives such as Rose Bengal and Erythrosin B identified from *in vitro* screens with truncated unregulated *E. coli* SecA displayed IC_50_ of 0.5 and 2 μM, respectively. Rose Bengal strongly inhibits preprotein translocation *in vitro* with IC_50_ of ~0.25 μM. (Huang et al., 2012b). Arylomycin, with a concentration < 4 μg/mL, inhibits the extracytoplasmic localisation of 98 proteins in *E. coli* (Walsh et al., 2019). (Pannomycin, identified in a differential sensitivity antisense assay (Parish et al., 2009), displays a non-validated inhibitory activity with an unclear mode of action. The N-(3-(benzyloxy)-5-ethoxybenzyl)-1-(piperidin-4-yl) methanamine (P87-A4) and its analog 2-((3-(benzyloxy)-5-ethoxybenzyl) amino)ethane-1-ol (17D9) were identified as inhibitors of SecA translocation ATPase by virtual ligand screening with IC_50_ values of 50.7 and 23.9 μM (De Waelheyns et al., 2015).

Other Sec pathway targets include signal peptidases that are attractive for antibiotics designing. Type I signal peptidase (SPase I) releases the mature secreted protein from the membrane-embedded SecY channel (Fig. 1A, yellow), and its activity is essential for cell growth and viability (Rao C.V et al., 2014). SPase I has a unique catalytic mechanism which makes it a target for inhibitors without affecting other eukaryotic serine proteases (Rao C.V et al., 2014). These inhibitors include monocyclic azetidinones that were effective against *E. coli* SPase I at a high concentration (500 μM) (Kuo et al., 1994); (5*S*)-Tricyclic penem that showed an IC_50_ of 0.2 and 5 μM against *E. coli* SPase I and methicillin-resistant *Staphylococcus aureus* SPase I(SpsB), respectively (Hu et al., 2003). Arylomycin C also inhibits *E. coli* SPase I potentially with IC_50_ 0.11–0.19 μM (Kulanthaivel et al., 2004).

As terminal Sec pathway components, the lipoprotein (Lol) and outer-membrane β-barrel proteins (OMPs) were targets for inhibitors. Some of the inhibitors of Lol pathway inhibit the chaperone LolA by preventing its binding to substrates (Choi and Lee, 2019). These inhibitors affect *E. coli* strains with MIC of 16 μg/ml (Ito et al., 2007; Pathania et al., 2009). Moreover, S-(4-chlorobenzyl)isothiourea and S-(3,4-dichlorobenzyl)isothiourea (A22) were effective against *E. coli* MG1655 LolA with IC_50_ of 150 and 200 μM, respectively (Barker et al., 2013). For OMPs, β-hairpin macrocyclic peptidomimetic JB-95 was described as an inhibitor for the BamA insertase and affects *E. coli* ATCC25922 with a MIC of 0.25 μg/ml (Urfer et al., 2016).

Here we took a more general approach aiming at inhibitors against the whole process of Sec-dependent post-translational protein secretion using *E. coli* as a screening model bacterium. To identify unknown new targets with potentially important secretory roles, we followed periplasmic secretion of alkaline phosphatase (PhoA) *in vivo*. PhoA is only active in this sub-cellular location after dimerization, and disulfide oxidation (Wanner et al., 1988; Manoil et al., 1990; Derman and Beckwith, 1991; Kriakov et al., 2003) and its activity was monitored using a HTS luminescence-based assay that we developed. Using this assay, a 240K small molecule library was screened. After hit confirmation and selection of analogs, fourteen compounds (HSI; HTS secretion inhibitors) belonging to 8 different chemical series with IC_50_’s ranging from 3 to 60 μM as determined in the secretion of alkaline phosphatase assay. The compounds were also tested as inhibitors of viability of either Gram-negatives or positives, including of 16 bacterial species (Gram^+^ and Gram^−^) from the WHO top pathogens list (Tacconelli et al., 2018).

Seven of the secretion inhibitor compounds, had microbicidal activity and represented three structural families HSI#9(parent) and 6; HSI#1(parent) and 5, 10 and HSI#12, 14. HSI#6, inhibited the growth of 8 Gram^**-**^ and 5 Gram^**+**^ bacteria with excellent IC_50_ values of 0.4-9 μM, while HSI#9 showed inhibition in the growth of 4 Gram^+^ strains and *E. coli* strains with IC_50_ <27 μM. Three compounds (HSI#1, 5 and 10) revealed antibacterial activity toward Gram^**+**^ (IC_50_ of ~7-38 μM) and of *E. coli* strain BW25113Δ*tolC* (IC_50_ <14 μM), which suggested that outer membranes hampered their permeability. HSI#12 and 14 inhibited only the growth of *E. coli* strains (IC_50_ of 44-65 μM). Moreover, HSI#9 prevented an EPEC strain from infecting HeLa cells. Under our assay conditions, none of the tested compounds inhibited the SecA ATPase activities *in vitro*; therefore the observed inhibition is not SecA-specific but rather targets additional unknown essential components of the post-translational secretion process.

In summary, our *in vivo* screening approach using an *E. coli* lab strain returned a wide range of useful anti-bacterials with broad and narrow spectrum properties.

## Results

### Development of an in vivo HTS assay for E.coli protein secretion

For the HTS we used *E.coli* strain BL21 expressing and secreting alkaline phosphatase (PhoA). Measurement of PhoA activity was used to monitor post-translational protein secretion. PhoA becomes enzymatically active only once translocated to the periplasm via a functional Sec machinery. In the presence of Sec pathway inhibitors, PhoA would not be translocated and phosphatase activity should be abrogated. We developed a sensitive 384 well setup phosphatase assay amenable to high throughput screening using “AP-juice” as a substrate (P.j.K GmbH; see Materials and methods) that can be monitored in a luminometer once hydrolyzed (Fig. 1B). The *prophoA* gene was expressed behind IPTG-inducible T7 RNA polymerase control on plasmid pIMBB882. The amounts of IPTG and number of cells used were optimized for the highest signal to noise ratio at 30 °C. As a positive control (i.e. maximal inhibition) we used the known inhibitor sodium azide (Huang et al., 2012a), cells treated with the DMSO vehicle alone served as a negative control.

### HTS results

To identify inhibitors of the post-translational Sec preprotein export pathway we tested a small molecule library of 238,601 compounds using the *in vivo* PhoA assay. The compounds were dissolved in 100% DMSO at 20-30 mM and tested at a final concentration of 20 μM (Lichstein and Soule, 1944). Under our HTS assay conditions, sodium azide inhibited the secreted phosphatase activity by >90 %. 1984 compounds out of the entire library inhibited the PhoA activity by >37.6 % relative to the positive (sodium azide) and negative control (DMSO). These were next tested in a dose-dependent manner up to 60 μM. The dose-response testing yielded 8 molecules representing 8 structural families that showed dose-dependent inhibition of *in vivo* PhoA secretion. 191 analogs of these 8 hits (resupplied for indepedent confirmation) were selected and tested in the *in vivo* PhoA secretion assay leading to the identification of a total of 14 compounds representing 8 families with dose-dependent inhibition (Fig. 2). None of these inhibited the enzymatic activity of purified native phosphatase in a counter assay.

**Figure 2.**
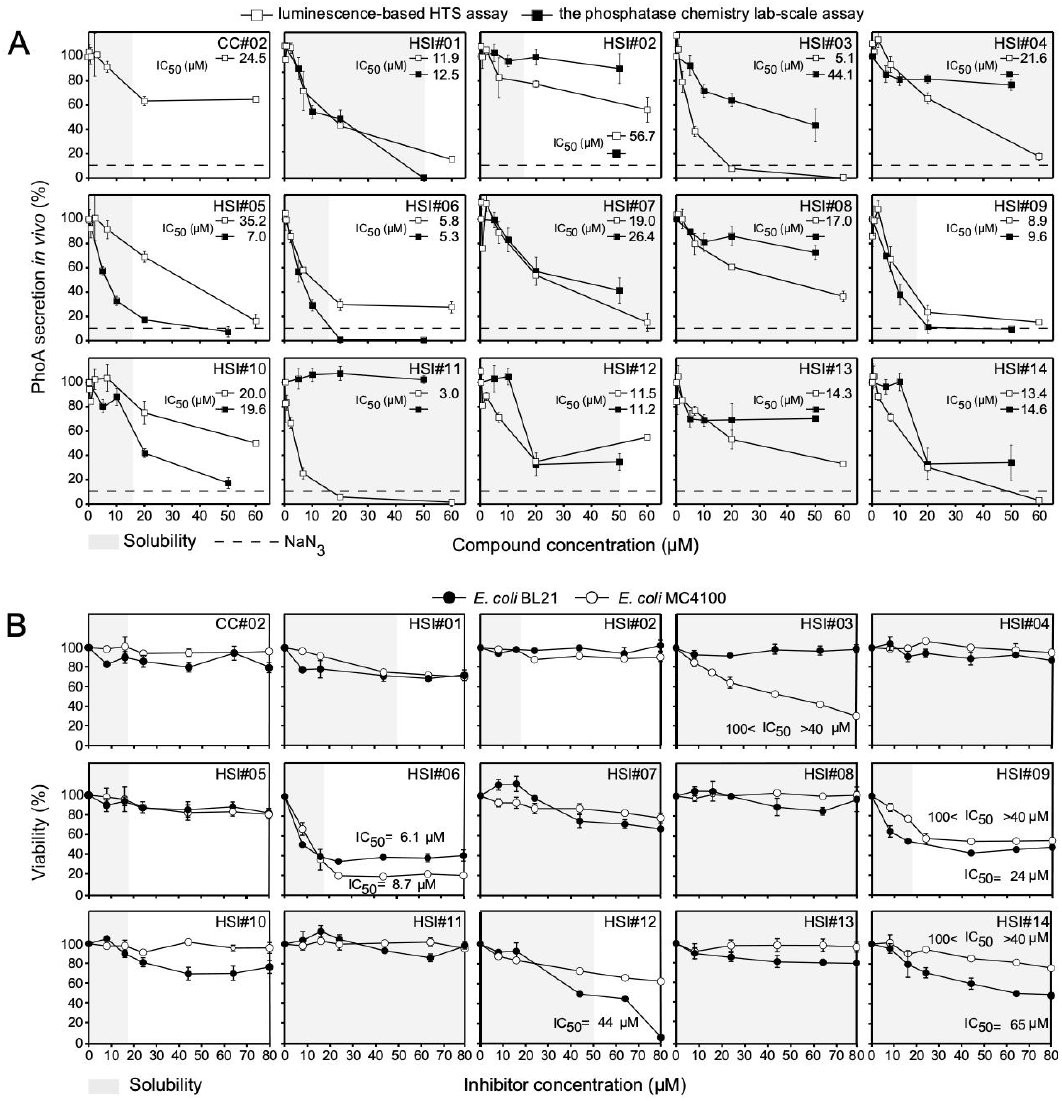
*In vivo* anti-PhoA secretion and anti-bacterial activity of secretion inhibitors toward *E. coli* strains. **A.** The effect of the 14 compounds discovered by HTS and CC#02 at different concentrations on PhoA secretion *in vivo* tested using the luminescence (HTS) and the *p*-nitrophenyl (lab-based) PhoA actvity assays. PhoA secretion in the absence of any compound but in the presence of DMSO 2.5% (v/v) was set as 100%. **B.** Inhibition of bacterial viability by the indicated inhibitors and CC#02. The growth of the indicated *E. coli* strains (OD_600_ in the absence of any compound but in the presence of 2.5% (v/v) dimethyl sulfoxide was taken as 100% and OD_600_ in the presence of inhibitor was normalized to it) was plotted against the inhibitor concentration. IC_50_ values for growth inhibition are indicated. *n=3*. The results are presented as the mean ± SD. Gray shade: aqueous solubility.

The 14 HSI compounds of interest were re-purchased and their inhibitory activity re-confirmed in the luminescence-based *in vivo* secretion assay. All inhibited PhoA secretion (IC_50_<57 μM; Fig. 2; Table 1). Three of them very significantly (HSI#3, 6 and 11; IC_50_ of 3-5.8 μM; Fig. 2A; Table 1), 9 of them and CC#02 significantly (IC_50_ of 8.9-24.5 μM) and two weakly (HSI#5 and 2; IC_50_ of 32 and 56.7 μM, respectively). 10 of the compounds displayed very similar (HSI#1, 5, 6, 9 and 10) or similar (HSI#3, 4, 7, 12 and 14) trends in a lab-scale *in vivo* PhoA secretion assay (Gouridis et al., 2010; Jackson et al., 2016)(see Supplementary materials and methods) monitoring *p*-nitrophenyl phosphate production (Fig. 2A; Table 1), while the remaining 4 compounds showed marginal effects.

**Table 1.** Properties of PhoA secretion inhibitors returned from the HTS

| Parent and daughter molecules | Secretion Inhibitor | Inhibition of PhoA secretion IC<sub>50</sub> [μM] |  |  | Bacterial viability IC<sub>50</sub> [μM] |  |  |  | Toxicity HEK 293 LD<sub>50</sub> [μM] | Aqueous solubility |  |
| --- | --- | --- | --- | --- | --- | --- | --- | --- | --- | --- | --- |
|  |  | HTS | Lab-scale | Lab-scale (Δ*toI*C strain) | Gram<sup>-</sup> |  |  | Gram<sup>+</sup> |  | μM | μg/ml |
|  |  |  |  |  | *E. coli* |  |  | *B. subtilis* |  |  |  |
|  |  |  |  |  | BL21 | MC4100 | BW25113 |  |  |  |  |
|  | CC#01* | 237 | NA |  | 4 | 4 |  | >100 | nt | nt | nt |
|  | CC#02** | 24.5 | NM |  | NM | NM |  | 18.8 | 60.2 | 16.7 | 4.1 |
| Structure family 1 |  |  |  |  |  |  |  |  |  |  |  |
| HTS hit | HSI#03 | 5.1 | 44.1 | 21.0 | NM | 40-100 | NM | NM | 60.2 | 150 | 40.0 |
| Structure family 2 |  |  |  |  |  |  |  |  |  |  |  |
| HTS hit | HSI#07 | 19.0 | 26.4 | 19.9 | NM | NM | NM | NM | 60.2 | 150 | 32.4 |
| Analog | HSI#12 | 11.5 | 11.2 | 26.0 | 60 | NM | 39.7 | NM | 60.2 | 50.0 | 11.8 |
| Analog | HSI#14 | 13.4 | 14.6 | 11.5 | >100 | NM | 31.3 | NM | 60.2 | 150.0 | 33.5 |
| Structure family 3 |  |  |  |  |  |  |  |  |  |  |  |
| HTS hit | HSI#09 | 8.9 | 9.6 | 0.91 | 40-100 | 40-100 | 5.4 | 22.5 | 20.1 | 16.7 | 5.3 |
| Analog | HSI#06 | 5.8 | 5.3 | 0.33 | 6.1 | 8.7 | 26.4 | 2.4 | 20.1 | 16.7 | 5.7 |
| Structure family 4 |  |  |  |  |  |  |  |  |  |  |  |
| HTS hit | HSI#01 | 11.9 | 12.5 | 1.4 | NM | NM | NM | 37.6 | 6.7 | 50 | 12.5 |
| Analog | HSI#05 | 32.0 | 7 | 0.94 | NM | NM | NM | 27.8 | 6.7 | 16.7 | 4.8 |
| Analog | HSI#10 | 20.0 | 19.6 | 0.99 | NM | NM | NM | 26.0 | 6.7 | 16.7 | 4.4 |
| Structure family 5 |  |  |  |  |  |  |  |  |  |  |  |
| HTS hit | HSI#11 | 3.0 | NM | - | NM | NM | NM | NM | 60.2 | 150.0 | 23.4 |
| Structure family 6 |  |  |  |  |  |  |  |  |  |  |  |
| HTS hit | HSI#13 | 14.3 | NM | - | NM | NM | NM | NM | 60.2 | 150.0 | 57.7 |
| Analog | HSI#08 | 17.0 | NM | 32.6 | NM | NM | NM | NM | 60.2 | 150.0 | 54.4 |
| Structure family 7 |  |  |  |  |  |  |  |  |  |  |  |
| HTS hit | HSI#04 | 21.6 | >50 | 7.8 | NM | NM | NM | NM | 60.2 | 150.0 | 37.6 |
| Structure family 8 |  |  |  |  |  |  |  |  |  |  |  |
| HTS hit | HSI#02 | 56.7 | NM | 14.9 | NM | NM | NM | NM | 20.1 | 16.7 | 5.2 |

CC : Control compound
\* : NaN<sub>3</sub>; concentration in (mM); value indicates MIC not IC<sub>50</sub>\*\* : Compound is PubChem ID 11528894, proposed as a low micromolar inhibitor of *E.coli* (Crowther et al., 2015)
nt : Not tested.
NA : Not applicable
NM : Non-measurable

None of the compounds directly inhibited SecA ATPase activities *in vitro* (Fig. S1 and 2) and thus the inhibition of PhoA secretion seen resulted from an effect on a different target.

Next, the HSI compounds were examined for solubility and cytotoxicity and for microbicidal activity against several species and strains.

### Solubility and cytotoxicity testing of the HSI compounds

The kinetic aqueous solubility of the 14 compounds was determined using a turbidimetric method, measuring an increase in the absorbance (at 570 nm) of scattered light resulting from compound precipitation. CC#02, HSI#5 and 10 displayed low solubility, (4.1-4.8 μg/ml; 16.7 μM) (Table 1), close to the minimal solubility recommended for drugs (U.S. Pharmacopeia, (Savjani et al., 2012; Lipinski, 2002). HSI#2, 6 and 9 showed solubility just over 5 μg/ml (16.7 μM) while the remaining nine compounds showed higher solubility (10-60 μg/ml; 50-150 μM) (Table 1).

The cytotoxicity of the compounds was tested toward HEK293T cells by determining the number of remaining cells with an ATP monitoring system (see Materials and methods; 1016739, PerkinElmer). HSI#1, 5 and 10 showed the highest toxicity levels (LD50=6.7 μM); this likely compromises their use as lead structures for antimicrobial development (Table 1).

### Effect of HSI compounds on the viability of Gram^−^ bacteria

We next determined the *in vivo* antimicrobial properties against both Gram^**-**^ (Fig. 2B) bacteria for the 14 compounds and the controls.

Sodium azide inhibited growth by ~80% at 4 mM, while CC#02 barely inhibited growth of any of the *E. coli* strains. In the absence of any other relevant indicator of secretion inhibition, sodium azide sets a boundary of what level of anticipated inhibition of secretion might be lethal for ~80% of the cells. At the concentrations used, sodium azide might have pleiotropic effects.

Viability of *E. coli* strains BL21 and MC4100 was inhibited significantly by HSI#6 (IC_50_ of 6.1-8.7 μM; Fig. 2B; Table 1) and more weakly from HSI#9, 12 and 14 (24-100 μM; Fig. 2B; Table 1). On the contrary, none of the 14 compounds affected the growth of Enteropathogenic *E. coli* (Fig. S3).

### Effect of the compounds on the viability of Gram^+^ bacteria and the WHO top critical pathogens

We next tested the compounds on the viability of Gram^+^ bacteria: a non-pathogenic *B. subtilis* lab strain and four strains closely related to the Gram^+^ ones from the WHO top 16 critical list: *S. aureus* ATCC 6538P, *Enterococcus faecalis* ATCC 804B, *Streptococcus pneumoniae* and *Mycobacterium abscessus* ATCC19977 (Table 1 and 2). Sodium azide barely inhibited the growth of most Gram^+^ bacteria (any observable inhibition required commonly >100 mM; Table 1). CC#02 inhibited three of the Gram^+^ bacteria well (IC_50_ of 13-28 μM) but not *M. abscessus* (Fig. 3A). As with Gram^−^ bacteria, HSI#6 showed the highest antibacterial effect toward Gram^+^ bacteria (IC_50_ of 0.4-2.4 μM) (Fig. 3D; Table 1 and 2), with *S. pneumoniae* 2-3 times more sensitive. Moreover, HSI#1, 5, 9 and 10 that displayed limited effect against Gram^+^ bacteria, showed high antibacterial activity toward *B. subtilis, S. aureus*, *M. abscessus* and *E. faecalis* (IC_50_ of ~4-38 μM) (Fig. 3B, C, E and F; Fig. S4). The IC_50_ values of HSI#1 and 10 were, respectively, higher for *S. pneumoniae* compared to other Gram^+^ bacteria.

**Table 2.** Priority pathogens list for R&D of new antibiotics. Adjusted from the WHO 2018 recommendation list (Tacconelli et al., 2018)

| Pathogen list | Pathogen used in this study | Bacterial viability (IC<sub>50</sub>, µM) |  |  |  |  |  |
| --- | --- | --- | --- | --- | --- | --- | --- |
|  |  | CC#02** | HSI#01 | HSI#05 | HSI#10 | HSI#09 | HSI#06 |
| **Priority 1: Critical** |  |  |  |  |  |  |  |
| Multidrug-resistant and extensively-resistant *Mycobacterium tuberculosis* |  |  |  |  |  |  |  |
| *Mycobacterium tuberculosis* | *Mycobacterium abscessus* ATCC19977 | NM | 33.8 | 17.9 | 21.8 | 43.5 | 1.1 |
| *Pseudomonas aeruginosa*, carbapenem-resistant |  |  |  |  |  |  |  |
| *Pseudomonas aeruginosa* | *Pseudomonas aeruginosa* 3/88 | NM | NM | NM | NM | NM | 6.3 |
| *Enterobacteriaceae*, carbapenem-resistant, 3rd generation cephalosporin-resistant |  |  |  |  |  |  |  |
| *Klebsiella pneumoniae* | *Klebsiella pneumoniae* ATCC 27799 | NM | 80-100 | NM | NM | 80-100 | 8.7 |
| Enteropathogenic *E. coli* | Enteropathogenic *E. coli* O127:H6 strain E2348/69 | NM | NM | NM | NM | NM | >90 |
| *Enterobacter cloacae* | *Enterobacter cloacae* | NM | NM | NM | NM | 50 | 7.8 |
| *Proteus vulgaris* | *Proteus vulgaris* | NM | 80-100 | NM | NM | 34 | 2.9 |
| *Providencia stuartii* | *Providencia stuartii* | NM | NM | NM | NM | NM | NM |
| *Morganella morganii* | *Morganella morganii* | NM | NM | NM | NM | 70 | 6.4 |
| *Serratia marcescens* | *Serratia marcescens* | NM | NM | NM | NM | NM | 22.7 |
| **Priority 2: High** |  |  |  |  |  |  |  |
| *Enterococcus faecium* | *Enterococcus faecium* ATCC 804B | 28.0 | 34.5 | 9.4 | 23.5 | 6.9 | 0.9 |
| *Staphylococcus aureus* | *Staphylococcus aureus* ATC6538P | 13.8 | 14.8 | 4.1 | 7.8 | 6.0 | 1.0 |
| *Campylobacter spp* | *Campylobacter jejuni* | NM | NM | NM | NM | NM | 64.2 |
| *Salmonella spp* | *Salmonella typhimurium* | NM | NM | NM | NM | 80-100 | 12.4 |
| **Priority 3: Medium** |  |  |  |  |  |  |  |
| *Streptococcus pneumoniae* | *Streptococcus pneumoniae* | 113.3 | 77.8 | 29.7 | 60.9 | 12.6 | 0.4 |
| *Haemophilus influenzae* | *Haemophilus influenzae* | NM | NM | NM | NM | NM | 50.4 |
| *Shigella spp* | *Shigella sonnei* | NM | NM | NM | NM | 50 | 6.2 |

\*\* : This compound is PubChem ID 11528894; proposed as a low micromolar inhibitor of *E.coli* secretion (Crowther et al., 2015)
NM : Non-measurable

**Figure 3.**
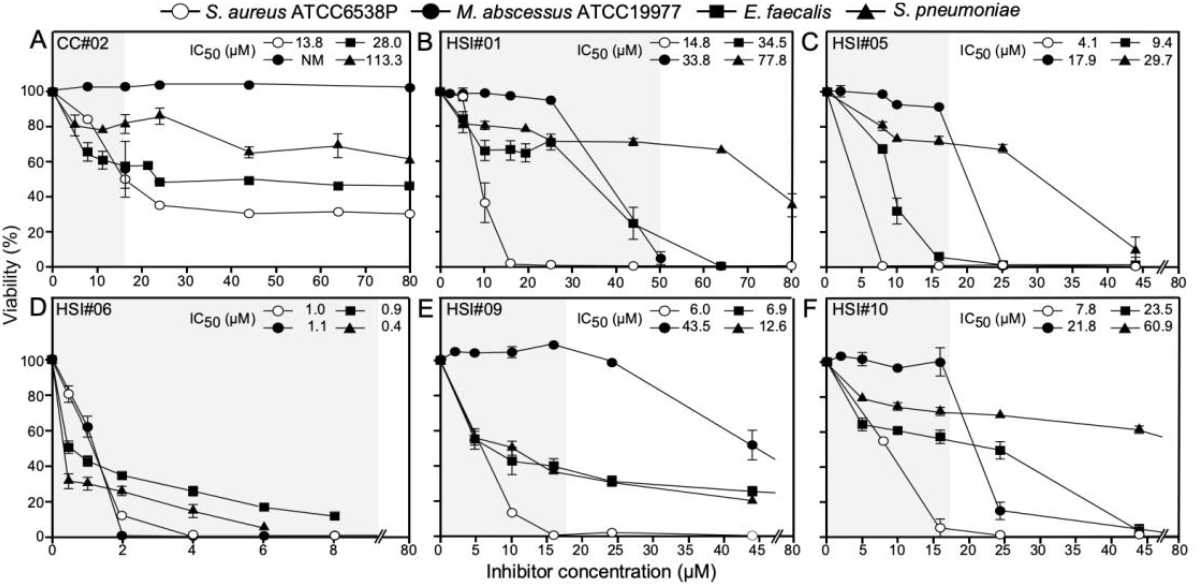
Antibacterial activity of the secretion inhibitors toward Gram^+^ bacteria. **A.-F.** Inhibition of bacterial viability by the indicated inhibitors and CC#02. Growth of the indicated Gram^+^ strains was plotted against the inhibitor concentration (as in Fig. 2B). IC_50_ values are indicated. *n=3*. The results are presented as the mean ± SD. Gray shade: aqueous solubility.

HSI#6 that inhibited the viability of *E. coli* and the five Gram^+^ bacteria, also inhibited the viability of 8 of the 12 Gram^−^ bacterial strains of the WHO top 16 critical list (Tacconelli et al., 2018)(Fig. 4; Table 2) with IC_50_ values of ~3-22 μM (Fig. 4A-D, F, G, J and K; Table 2).

**Figure 4.**
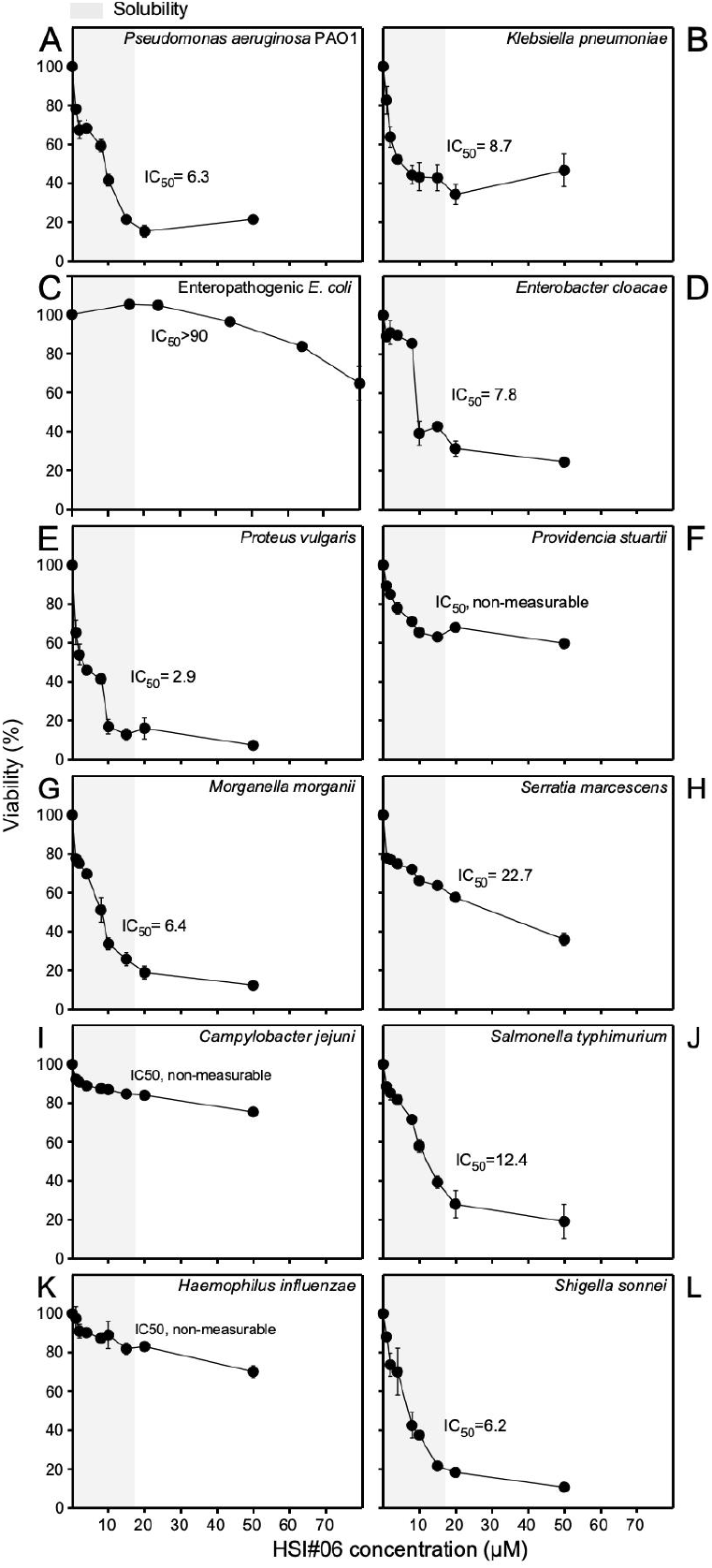
Antibacterial activity of secretion inhibitors toward 12 Gram^−^bacteria from the WHO top 16 list. **A.-L.** The indicated inhibitors which revealed inhibition of Gram^−^ bacterial viability. The growth of the indicated bacterial strains (As in Fig. 2B) was plotted against the inhibitor concentration. IC_50_ values for growth inhibition of the bacteria are indicated. Growth in the presence of 2.5% (v/v) dimethyl sulfoxide in the absence of inhibitors was taken as 100%. *n=3*. The results are presented as the mean ± SD. Gray shade: aqueous solubility.

### Effect of HSI compounds on the viability and secretion of E.coli strains with compromised outer membranes

The outer-membranes of Gram^−^ bacteria are a significant obstacle to novel antibacterial compound discovery (Balibar and Grabowicz, 2016). The outer membranes of *E.coli* strains *BW25113::imp-2413^+^* (Sampson et al., 1989) and *BW25113ΔtolC* (Baba et al., 2006) mutated or missing the outer membrane proteins LptD and TolC, respectively (Feriancikova et al., 2013; Zha et al., 2016), show increased permeability (Balibar and Grabowicz, 2016).

The *BW25113ΔtolC* and *BW25113imp-2413^+^* derivatives showed higher sensitivity toward 7 of the compounds (HSI#1, 5, 6, 9, 10, 12 and 14) by 2-40 times compared to BW25113, with *ΔtolC* showing in most cases stronger effects (Fig. 5A). Outer-membrane crossing reduced maximal protein secretion inhibition with 9 compounds inhibiting PhoA secretion in BW25113*ΔtolC* with IC_50_ lower than in BW25113 (HSI#1-6 and 8-10; Fig. 5B; Table 1). HSI#1, 5, 6, 9 and 10 inhibited PhoA secretion in BW25113*ΔtolC* with IC_50_ lower than that of the WT by 90%. HSI#2, 4 and 8 inhibited PhoA secretion only in BW25113*ΔtolC*.

**Figure 5.**
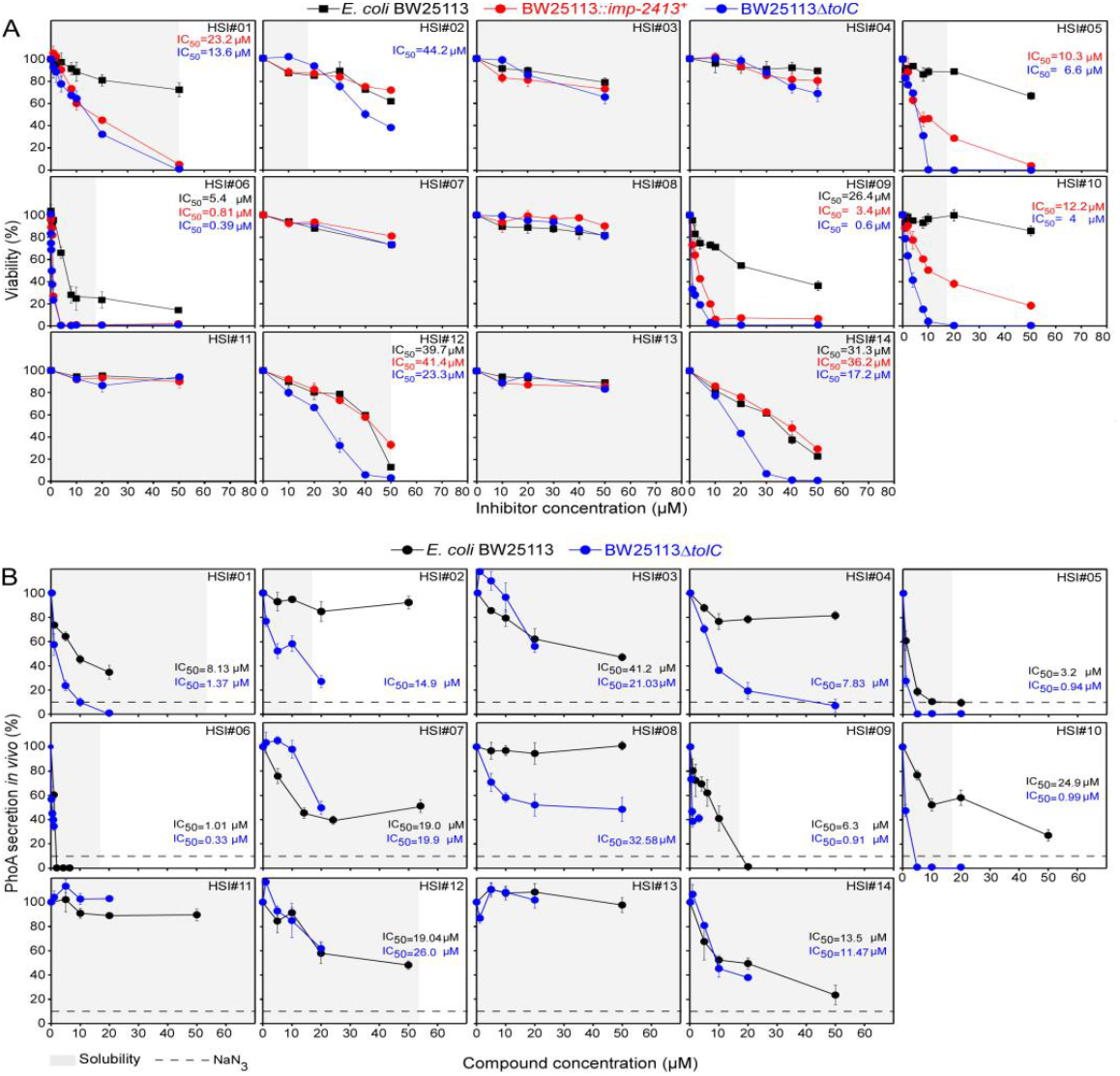
Antibacterial activity and *in vivo* PhoA secretion inhibition toward *E. coli* strains and derivatives. **A.** Antibacterial activity toward *E. coli* BW25113 and its derivatives BW25113*ΔtolC* and *BW25113::imp-2413^+^* of the indicated 14 inhibitors isolated from the HTS screening. *n=3*. Results are presented as the mean ± SEM. Gray shade: aqueous solubility. **B.** The effect of the 14 compounds isolated from the HTS screening on PhoA secretion *in vivo* of the indicated strains tested using the p-nitrophenyl assay (as in Fig. 2A). The SecA inhibitor sodium azide (4 mM) (Oliver et al., 1990) was used as a positive inhibitory control for maximal SecA inhibition observable *in vivo* (dashed line). Gray shade: aqueous solubility.

Apparently, for some compounds, outer membrane permeability was an obstacle in reaching sufficient concentrations in the cell to be inhibitory.

### Effect of compounds on virulence of enteropathogenic *E. coli*

The Sec pathway is important for the assembly of most cell envelope structures and other pathogenicity-specific protein export systems, like the Type 3 Secretion System (T3SS)(Portaliou et al., 2017), that are not essential for viability. Of the 11 non-cytotoxic compounds HSI#9 reduced virulence of EPEC has anti-virulence activity on the T3SS, as it inhibited infection of mammalian cells by EPEC at a concentration of 16 μM (Fig. S6) (Portaliou et al., 2016; Portaliou et al., 2017). It inhibited EPEC colonization of HeLa cells and allowed neither actin pedestals nor EspA fillaments to form (Goosney et al., 2000)(Fig. S6).

### Chemical characterization of derived PhoA secretion inhibitors

HSI#1(parent), 5 and 10 belong to the same chemical series (Fig. 6A). The three active analogs bear a 1,2,3-thiadiazole ring connected to a lipholilic aromatic moiety with an acrylate linker making them potential Michael acceptors. Early SAR data gathered from the commercial tested analogs showed that this acrylate linker seemed to be essential for activity (data not shown) but more work will have to be done to confirm that initial observation.

**Figure 6.**
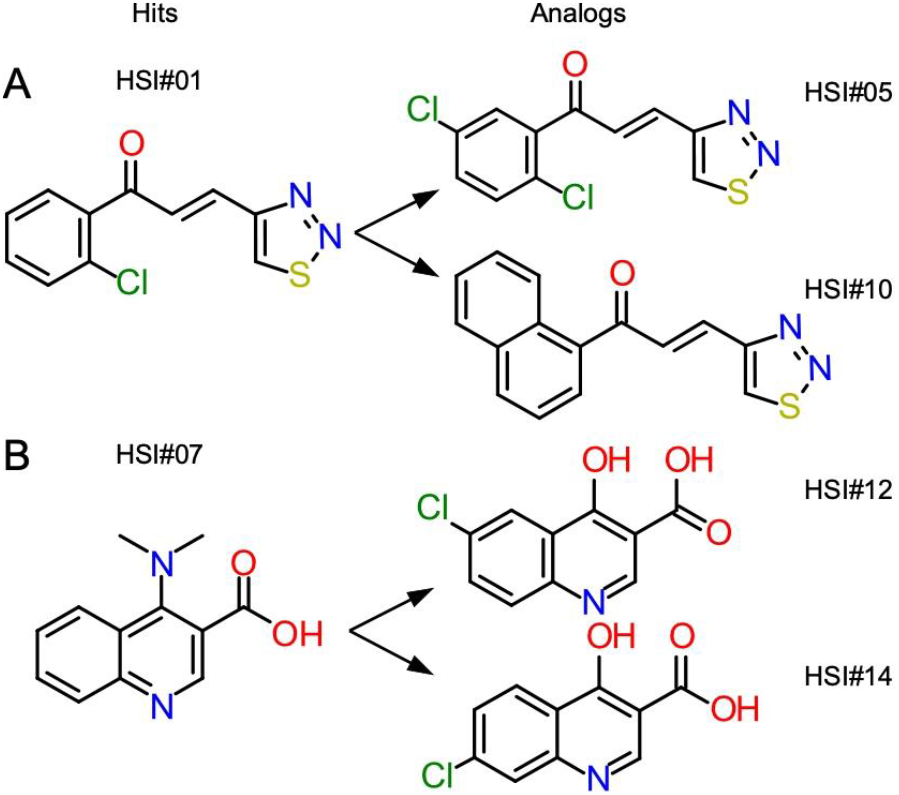
Compound structures of HSI#1, 5 and10 (A) and HSI #7, 12 and 14 (B).

HSI#7 (parent), 12 and 14 belong to the same chemical series (Fig. 6B), the three active analogs are derivatives of quinoline-3-carboxylic acid. Different derivatives of 3-quinolinecarboxylic acid have been reported to exhibit antimalarial (Coleman et al., 2001; Sharma et al., 2012) and antibacterial activities against both Gram^−^ and Gram^+^ (Wentland et al., 1984; Reuman et al., 1995).

HSI#6 and 9 will be characterized in-depth in a future study.

## Discussion

We present a multi-step pipeline to identify novel inhibitors of post-translational, bacterial protein secretion that display antibacterial activity (Fig. 1B). This pipeline returned 14 compounds from 8 structural families, that inhibited PhoA secretion with IC_50_’s <50 μM (Fig. 2A), and 7 of which showed antibacterial activity. Therefore, inhibition of protein secretion is validated as a tool to identify broad-spectrum antibiotics.

Five compounds (HSI# 1, 5, 10 and 6, 9) were effective both as secretion inhibitors (IC_50_ of ~5-35 μM) and as antibacterials (IC_50_ of ~1-37 μM) with HSI#6 being the most effective inhibitor toward both Gram^+^ and Gram^−^ bacteria. IC_50_ values are comparable with those of commercially available antibiotics, i.e. vancomycin, penicillin G, and ampicillin (IC_50_=17.0, 0.5, 0.01 μM toward *S. aureus*, respectively; Fig. S5). Given its only moderate toxicity on human cells (Table 1) and relatively broad-spectrum, HSI#6 may be a good lead for further optimization.

Three compounds (HSI#1, 5 and 10) displayed significant activity against Gram^**+**^ bacteria (Fig. 3A). This revealed that although a Gram^**-**^ bacterial model strain was used for screening, our approach can return potent Gram^+^ antibacterials. The screening assay is sensitive enough to pick out multiple, broad secretion inhibitors because it likely inhibits a universal process. These compounds reduced PhoA secretion in *E. coli* (Fig. 2A) but did not affect viability (Fig. 2B) suggesting inefficient penetration through the Gram^**-**^ outer membrane. This is common for many antibiotics that are highly effective against Gram^**+**^ bacteria (e.g. macrolides, novobiocin, rifamycin, lincomycin, clindamycin and fusidic acid; (Nikaido, 2003)). *S*trains missing outer membrane proteins (Augustus et al., 2004) increased the sensitivity of *E. coli* to 8 compounds including the three Gram^+^ antibacterials (HSI#1, 5 and 10) and the previously non-effective compound HSI#2 (Figure 5A). Additionally, the *tolC* knock out enhanced the potency of 9 compounds to inhibit PhoA secretion *in vivo* by 90% (Figure 5B), which would suggest using the *tolC* mutant as a tool for broad antibacterials selection. As HSI#1, 5 and 10 display high-level HEK293 toxicity, they are not attractive for further optimization, while HSI#9 may be a possible lead for optimization of Gram^+^-specific antibiotics. Although HSI#9 had only moderate to low antimicrobial activity against Gram^−^ bacteria, it inhibited EPEC virulence, suggesting that it might render secreted pathogenicity factors limiting for infection.

Nine of the remaining compounds were moderate to strong PhoA secretion inhibitors. Two compounds (HSI#12 and 14) affected only the growth of *E. coli* with IC_50_ >45 μM. As protein secretion is an essential process, we consider three possibilities for these false positives. Some HSI compounds might: (i) affect folding and/or formation of disulfides of periplasmic PhoA polypeptides that have already been secreted or are being secreted. Enzymes like the Dsb proteins are known catalysts of disulfide oxidation but are not essential for viability (Goodall et al., 2018; Loos et al., 2019). (ii) compromise the secretion process directly but not sufficiently so as to yield a substantial anti-bacterial effect. It should be noted that even sodium azide, a potent *E. coli* anti-bacterial at 3-4.6 mM (Lichstein and Soule, 1944), still yields a substantial level of secreted PhoA (~17%; Fig. 2A). Therefore, we anticipate that unless secretion is inhibited at such levels, it will not lead to lethality. In most cases the best correlation with sensitivity to a drug was their ability to be taken up at sufficient final intra-cytoplasmic concentrations (Liu et al., 2019). For some compounds maximal usable amounts are limited by their solubility (Table 1). For many essential cellular targets even reduction in production by 97% does not lead to lethality (Cui et al., 2017a) (iii) As the assay was run against *E.coli*, it remains possible that secretory inhibition may involve components that are not operational in Gram positive bacteria.

HSI#7, 12 and 14 are quinoline-3-carboxylic acid derivatives (Fig. 6B) and might inhibit DNA gyrase A (Domagala et al., 1986; Tunitskaya et al., 2011) as do analogous compounds (Reuman et al., 1995). How these activities might connect to protein secretion is unclear but it is known that such compounds can affect the expression of more than 100 genes in *Streptococcus* and induce oxidative stress (Chu et al., 2019). Similarly, although thiadiazole ring compounds have a broad spectrum of pharmacological activities including as anti-inflammatory, antiviral and antibacterial agents (Jain et al., 2013), it is not currently known how the 1,2,3-thiadiazole ring compounds (HSI#1, 5 and 10) (Fig. 6A) might affect protein secretion. Interestingly, thiouracil derivatives containing a triazolo-thiadiazole moiety have been developed and proposed to act as SecA inhibitors (Huang et al., 2012c; Cui et al., 2013; Cui et al., 2017b).

In summary, these results validated our new HTS approach by yielding starting points for potential new drug development. Future focusing of screening efforts to different reporter enzymes with different degrees of essentiality and topologies is expected to expand the gamut of promising compounds returned by this approach.

## Materials and methods

### Small compound library

The library was provided by HDC and contains around 240’000 drug-like and lead-like compounds, carefully selected by a team of medicinal and computational chemists to provide the best chemical starting points for drug discovery. The collection consists of three subsets (Discovery set, Explorer set and Probe set) which taken together provide an excellent diversity for drug discovery projects. More information can be found here: **https://www.hit-discovery.com/services/**

### HTS assay for bacterial proPhoA secretion *in vivo*

The *E. coli* strain BL21 (pLysS) with proPhoA (pIMBB882) was grown overnight in 5 ml LB containing Ampicilline (100 μg/ml) and Chloramphenicol (25 μg/ml) by taking a stab of the frozen glycerol stock and suspending it in the LB medium at 37°C. This was shaken at 250 rpm. The next day, 5 ml of the pre-culture was transferred into 200 ml LB medium and grown for another 2.5 hours at identical conditions until the OD_600_ reached 0.6. 25 μl of this cell suspension was dispensed into, full white 384 well, assay-ready plates containing 300nl from 2mM stock of the compounds (final concentration 20 μM). The plates were shaked vigorously for 30 sec and incubated at 30°C for 15 min. After this pre-incubation the *phoA* expression was induced by the addition of 5 μl (0.1 mM) IPTG followed by incubation (1.5 h; 30°C). Next, 5 μl of CellLytic express (Sigma-Aldrich) was added, shaked for 30 sec, and incubated at 20°C for 15 min. PhoA was detected by the addition of 25 μl of AP-juice (p.j.k. GmbH, Kleinblittersdorf, Germany), shaked for 60 sec and incubated (10-20 min), followed by Luminescence measurement on an Envision luminometer (Perkin Elmer).

### HTS screening

All compounds were dissolved in DMSO (20 mM stock) and were tested at a final concentration of 20 μM in 384 wells plates containing 320 compounds, 32 negative controls (DMSO alone) and 32 positive controls (Sodium Azide 4 mM). Controls were used to calculate for every compound the percent inhibition relative to the controls, as well controls are used to define the quality of the experiment per plate by calculation of the Z’-score and Signal over Background ratio (S/B). All plates screened had a Z’-score higher than 0.5 with an average of 0.78, and an average S/B of 25.8.

To exclude molecules that are giving false-positive results we developed a counter screen that was based on the same detection methodology as the primary PhoA screening assay. For this, 300 nl of compound/vehicle was spotted into a white 384 well plate, per well with a solution of Alkaline Phosphatase (EF0651, Thermo Scientific) in PBS (with Ca^2+^ and Mg^2+^) was dispensed (30 μl) and shaked. After 1-5 min incubation, 25 μl of AP-juice was added to the samples followed by a shaking step for 1 min and finally read on an Envision. Any compound that inhibited the PhoA activity in a dose-dependent manner was excluded from further evaluation.

### Solubility assay

The determination of the aqueous solubility of compounds in this assay is based on the principle of turbidimetry. Turbidimetric methods rely on the measurement of light scattering from precipitate in solution to determine the solubility. Precipitation is identified by an absorbance increase due to blockage of the light by the particles at the wavelength of 570 nm. Compounds, stored in matrix vials (Thermo Fisher) at a stock concentration of 30 mM or 10 mM in 100% Dimethyl sulfoxide (DMSO), are used to make a serial dilution (dose-response) in a 96-well v-bottom propylene plate (Greiner, 651201). Serial dilutions are made to perform the solubility assay, they are made row-wise and start with undiluted compound (30 mM or 10 mM) in the first well and are then 1 over 3 further on diluted. The serial dilutions are 8 doses long and contain 6 compounds per plate maximal (rows B-G). Columns 1 and 12 are filled with 100% DMSO for control purposes. The dose-response plates are 200 times diluted in 300μL Phosphate Buffered Saline (PBS; pH= 7-7.2 without Ca/Mg) (Gibco, 14190-144) by transferring 1.5 μL of serially diluted compound. This results in final starting concentration of 150 μM (starting from 30 mM) or 50 μM (starting from 10 mM) at 0.5% DMSO. Dilutions are made by diluting one 96-well plate in two 96-well, flat-bottom, polystyrene plates (Greiner, 655101). These plates are then incubated at room temperature for 1h. After 1h incubation, the plates are read on the envision (Perkin Elmer) at 570 nm.

The data of this assay is reported as “Soluble at” value for each compound. This “soluble at” value represents the concentration where the compound is still soluble and is the concentration before the first precipitated concentration. The first precipitated concentration is the concentration where the absorbance value is more than 5, standard deviations higher than the average background absorbance. The average background absorbance and the standard deviation are calculated on the 0.5% DMSO controls in columns 1 and 12.

### Cytotoxicity assay

Hek293T cells which are harvested and diluted obtaining a cell suspension with a concentration of 200.000 cells/ml in complete growth medium (DMEM, 4.5 g/l d-glucose, pyruvate 1 mM, 0.075% bicarbonate and 10% Fetal Bovine serum). Of this cell suspension 50 μl (10.000/well) is seeded in a 384 wells culture plate (white polystyrene, tissue culture treated) and incubated at 37° - 5% CO_2_ for 4 hours in a moisturized incubator which enables the cells to adhere. After the incubation small chemical compounds are diluted in medium and 10 μl of a compound solution is added onto the cells ending up with the required compound concentration. The cells are incubated for 24 hours at 37° - 5% CO_2_ in a moisturized incubator where after x μl medium is removed to equalize the liquid levels within one plate ending up with ±25 μl/ well. The amount of remaining cells/well is determined with ATPlite™ 1step (Perkin Elmer) AdenosineTriPhosphate (ATP) monitoring system which is based on firefly luciferase.

Cytotoxicity was calculated by subtracting RLU (Relative Light Units) obtained of the cells incubated with a compound from the RLUs obtained from cells in presence of vehicle.

The cytotoxic effect of a test compound was determined as: Percent cell death = [1-((RLU determined for sample with test compound present – 1) divided by (RLU determined in the presence of vehicle – 1))] * 100

### Antimicrobial activity test

The antibacterial activity of compounds against various bacterial strains (*S. aureus* ATC6538P, *B. subtilis* ATCC6633, *E. coli* BL21, *E. coli* MC4100, *E. coli* BW25113, *E. coli* BW25113Δ*tolC*, *E. coli* BW25113Δ*lptD* and Entheropathogenic *E. coli O127:H6 (strain E2348/69 / EPEC*)) was measured using the serial dilution method in microplates. An overnight culture of all tested bacteria in LB medium was diluted 200-fold in fresh LB medium and incubated at 37°C until the OD_600_ reached 0.3. Nine strains of the WHO top 16 list pathogens, *Pseudomonas aeruginosa* 3/88, *Klebsiella pneumoniae* ATCC 27799, *Enterobacter cloacae, Proteus vulgaris, Providencia stuartii, Morganella morganii*, *Serratia marcescens*, *Salmonella typhimurium* and *Shigella sonnei* were grown in LB medium, while *Mycobacterium abscessus* ATCC19977, *Enterococcus faecium* ATCC 804B, *Campylobacter jejuni, Streptococcus pneumoniae, Haemophilus influenzae* and *Mycobacterium abscessus* were grown in tryptic soya broth (TSB) and incubated at 37°C and 5 % CO_2_ until the OD_600_ reached 0.3 as previously discussed. Next, 20 μl of this culture, which was previously diluted to OD_600_<0.01, was added to a 96-well microtiter plate containing different concentrations of each compound in the range of 0-100 μM (final DMSO concentration 2.5% (v/v); final volume of 200 μl) or DMSO alone (2.5% (v/v)). Bacterial cultures were incubated at 37°C for 20 hours with no shaking, OD_600_ was measured spectrophotometrically (Tecan Infinite^®^ 200 PRO) and the data were normalized against a control culture [0 μM compound, 2.5% (v/v) DMSO]. IC_50_ values were calculated in GraphPad Prism by nonlinear regression using equation model: Y=Y_Bottom_+(YTop-Y_Bottom_)/(1+10^((Log IC_50_-X)*(−1.0)) where Y_Bottom_ and Y_Top_ are plateaus in the units of the Y axis. The IC_50_ gives a response halfway between Y_Bottom_ and Y_Top_ thus measure of the potency of a compound in inhibiting bacterial viability and indicates reduction of bacterial growth by 50%.

### Immunofluorescence microscopy to follow filamentation of EPEC and infection of HeLa cells

An EPEC cells grown in LB medium (15 h; 37°C) was diluted in 5 ml pre-warmed DMEM 100-fold with 16 μM concentration of each of compounds. Bacteria were then grown at 37°C and 5% CO_2_ in a stationary incubator (non-shaking conditions) until OD_600_ reached 0.6. The sub-confluent lawns of HeLa cells, which had grown on glass coverslips inside a 24-well tissue culture dish (approximately 40,000 cells per well), were infected for 2 hours with the primed bacterial cultures (~3.5×107 bacteria/in 500 μl inoculated in each well). After 2 hour infection, the non-adhered bacteria were removed with three sequential gentle washes with phosphate-buffered saline (PBS) solution. The coverslips underwent fixation (3% formaldehyde solution in PBS; 30 min; rt.), permeabilization (1% Triton X-100 in PBS; 4 min; rt.), blocking (1% BSA in PBS; 30 min; rt.) with intermediate washing steps (3 times with 1 × PBS). Next, the wells were incubated with α-EspA (500-fold dilution; Davids Biotechnologie antibody (1:20,000 dilution detects 100 ng pure EspA in 1:20,000 dilution after 10 minutes exposure with SuperSignal West Pico PLUS Chemiluminescent Substrate from Thermo Scientific); 30 min; rt.)(37). Afterwards, the cells were washed 3 times with 1×PBS and the EspA filaments were stained with a donkey α-rabbit secondary antibody (500-fold dilution in 1% (v/v) BSA solution in PBS; Cy™3 AffiniPure Donkey Anti-Rabbit IgG (H+L), code number 711-165-152, Jackson Immunoresearch) and in parallel, the HeLa cell actin was detected with phalloidin (500-fold dilution in 1% BSA solution in PBS; Oregon Green™ 488 Phalloidin, Catalogue Number O7466,Invitrogen) and both bacterial and eukaryotic DNAs were stained with 1/1000 TOP-RO-3 solution (1000-fold dilution, TO-PRO™-3 Iodide (642/661), Catalogue Number T3605, Thermo Fisher Scientific, MA, USA). The coverslips were mounted on slides using Prolong Gold anti-fade reagent (Invitrogen), dried (dark; 12h) and observed using an Axioplan 2 widefield fluorescence microscope (Carl Zeiss MicroImaging GmbH, Germany). The images were acquired using a Hamamatsu ORCA R2 camera and computer-processed using Axiovision software version 4.8.2.0 (Carl Zeiss MicroImaging GmbH, Germany). The pictures were processed using ImageJ software version 1.8.0 and colours were added artificially using Photoshop Software version CS6 (Savjani et al., 2012; Portaliou et al., 2017; Lipinski, 2002).

## Acknowledgements

We are grateful to: Prof. F. Claessens (KU Leuven) for use of a Luminoskan Ascent Microplate Reader (Thermo); to HDC (https://www.hit-discovery.com/) for providing the compound library and for their input and help in the further optimization of the assay for use on the screening platform; to N.Eleftheriadis for discussions.

## Contribution statement

H.M.B. performed viability, secretion and ATPase assays; B.E. performed preliminary viability experiments and HeLa infections; A.L., M.A., K.H and C.P. organized the library, synthesized and characterized compounds and performed the HTS luminescence-based assay, cytotoxicity and solubility experiments; D.J. performed *in vitro* ATPase assays; L.M. and B.E. performed HeLa infection assays; A.J. analyzed data; E.A. and H.M.B wrote the paper and analyzed data; E.A. and K.S. conceived, managed and supervised the project. All authors read and approved the MS.

